# Perinatal exposure to individual and a metal mixture induces persistent and sex-specific alterations to the cardiac transcriptome of offspring

**DOI:** 10.64898/2026.07.11.737950

**Authors:** Morgan Steiner, Jason Laird, Sylvia S. Sanchez, Susmitha Pagadala, Shyam Biswal, Fenna C. M. Sille, Mark J. Kohr

**Author notes:** **Corresponding Author:** Mark J. Kohr, PhD, FAHA, FCVS, Department of Environmental Health and Engineering, Johns Hopkins Bloomberg School of Public Health, 615 N. Wolfe Street, Room E7616, Baltimore, MD 21205. These authors contributed equally to this study (Fenna C. M. Sille, Mark J. Kohr).

## Abstract

Exposure to heavy metals, such as lead, arsenic, cadmium, and chromium, has been individually linked to cardiac dysfunction during development and into adulthood. Although these metals are commonly encountered as a mixture, few studies have investigated the mixture effects of gestational exposure to these metals on the developing postnatal heart. To this end, we investigated the transcriptomic effects of individual heavy metals (arsenic, cadmium, chromium, and lead) and the combined mixture on female C57BI/6 mice prior to gestation through lactation. RNA was extracted from whole offspring hearts, and total RNA was sent for bulk RNA-sequencing. We found heavy metal exposure altered genes associated with circadian rhythm, cell division and DNA damage repair, and immune signaling. Moreover, we detected changes to the cellular composition of these hearts and an increase in *Il2ra* expression, indicating an increase in activated natural killer cells. When targeting postnatal heart development and maturation pathways, we found the mixture induced a general upregulation of almost all targeted pathways, which seemed to be driven by co-exposure to all metals instead of one metal driving the mixture phenotype, and revealed a potential functional-energetic mismatch. This study is one of the first to show that perinatal exposure to heavy metals altered circadian rhythm and immune signaling gene expression in a metal- and sex-specific manner, disrupted normal cardiac cellular composition, and upregulated genes associated with postnatal heart maturation.

## Introduction

Exposure to heavy metals and metalloids, including arsenic, cadmium, chromium, and lead, is of international importance due to environmental persistence and inability to be broken down via natural mechanisms. While these metals all occur naturally in the environment, an increase in environmental prevalence can be traced to anthropogenic processes, such as mining and smelting, pesticide use, and landfill leachate.^1–3^ Common routes of human exposure include ingestion of contaminated food and/or water, inhalation, and dermal exposure.^3^ Even exposure to low concentrations of these metals leads to adverse human health effects, underscoring why lead, arsenic, cadmium, and hexavalent chromium, are listed as 1^st^, 2^nd^, 3^rd^, and 30^th^, respectively, on the Agency for Toxic Substances and Disease Registry’s (ATSDR) substance priority list.

Arsenic, cadmium, and lead are well-characterized cardiotoxicants associated with cardiovascular disease in the general population,^4^ with emerging evidence suggesting that chromium also has a detrimental impact on the cardiovascular system.^5^ Moreover, prior work from our lab showed that acute exposure to arsenic increased susceptibility of the female mouse heart to ischemic injury,^6^ while chronic exposure increased blood pressure and altered cardiac structure in male mice.^7^ Furthermore, cadmium exposure had a detrimental impact on whole heart and isolated cardiomyocyte function in exposed male mice, but had no effect in females.^8^ Collectively, our findings highlight the sex-specific impact of metal exposure on the cardiovascular system in adult mice. In addition, previous studies from our lab have established the toxicity of metal exposure during critical periods of exposure, including pregnancy, as gestational arsenic exposure blunted maternal heart growth in pregnancy^9^ and increased maternal heart size postpartum.^10^ It is important to note that alterations to the maternal cardiovascular system can directly impact the developing fetus, and indeed we found that embryonic heart weight and body weight measurements were altered with gestational arsenic exposure.^9^ Additional studies link exposure to arsenic,^11^ cadmium,^12^ and lead^13^ to an increased risk of congenital heart defects. Arsenic and lead have also been linked to increased childhood blood pressure,^14,15^ while chromium may drive an inverse relationship between exposure and blood pressure in children,^16^ although additional reports have not detected an association between chromium exposure and childhood blood pressure.^17,18^ While the cardiotoxic effects of arsenic, cadmium, chromium, and lead have been characterized individually, few studies have investigated the effects of perinatal metal mixture exposure on the postnatal cardiac biology of offspring.

In the current study, we analyzed the effects of perinatal exposure to arsenic (in the form of sodium (meta) arsenite), cadmium (II), chromium (VI), and lead (II) individually and as part of a mixture at their respective regulatory limits set by the United States Environmental Protection Agency (EPA). Our previous work characterized the neurobehavioral effects of individual metals and the metal mixture,^19^ and herein, we find that perinatal exposure to metals alters the cardiac transcriptome in a metal- and sex-specific manner by interfering with genes associated with nuclear division, circadian rhythm, and immune signaling and altering postnatal heart maturation. This study highlights the importance of assessing heavy metal mixture exposures at environmentally relevant concentrations during critical periods of exposure, including pregnancy.

## Materials and Methods

### Mouse Exposure Model & Animal Care

Wild-type C57BL/6J mice (Jackson Laboratories, Bar Harbor, ME, U.S.A) were housed in the Johns Hopkins School of Public Health vivarium, which follows the Animal Welfare Act regulations and Public Health Service (PHS) Policy (PHS Animal Welfare Assurance #D16-00173 (A3272-01)) and is accredited by the Association for the Assessment and Accreditation of Laboratory Animal Care (AAALAC International # 000503). Our exposure model adheres to NIH guidelines for the care and use of laboratory animals and was approved by Johns Hopkins University’s Institutional Animal Care and Use Committee (Protocol # MO22H379), following the National Research Council’s Guide for the Care and Use of Laboratory Animals. Additionally, this study followed the ARRIVE 2.0 guidelines for reporting animal research^20^, where dams were randomly selected to metal treatment groups and assigned to animal cage identification numbers to blind researchers to treatment assignments during analysis.

Mice were maintained under a 12-hour reverse light-dark cycle and at 20-22°C and 55±5% humidity with *ad libitum* access to food and water. To reduce background metal exposure, mice were given purified, metal-free chow (AIN-93M; Research Diets, New Brunswick, NJ, USA, Cat# D10012Mi) and water (Crystal Geyser; CG Roxane, NY, USA). Food and water were replaced every 3 to 4 days and average intake was measured throughout the exposure period. Prior to breeding, males were housed individually in environmentally enriched cages (extra paper bedding and mouse igloo) and females were housed in groups of five per cage. Post-breeding, dams were housed individually in environmentally enriched cages.

Female mice were exposed to either individual metals or the mixture beginning two weeks prior to pre-timed mating and continuing through gestation and lactation. The following metal salts were used for exposure: lead (II) acetate trihydrate (Pb(C_2_H_3_O_2_)_2_·3H_2_O, CAS# 6080-56-4), sodium (meta) arsenite (NaAsO_2_, CAS# 7784-46-5), cadmium chloride (CdCl_2_, CAS# 10108-64-2), and sodium dichromate dihydrate (Na_2_Cr_2_O_7_·2H_2_O, CAS# 7789-12-0). Metals were dissolved in water: sodium (meta) arsenite (10 µg/L = 5.77 ppb of elemental arsenic), cadmium chloride (5 µg/L = 3.07 ppb of elemental cadmium), sodium dichromate dihydrate (100 µg/L = 34.9 ppb of elemental total chromium), and lead (II) acetate trihydrate (15 µg/L = 8.19 ppb of elemental lead), as previously described.^19^ These levels are well below the U.S. EPA’s defined maximum contaminant levels (MCLs - 10 µg/L total arsenic; 5 µg/L total cadmium; 100 µg/L total chromium)^21^ or the 2024-updated action level for lead, (AL, 10 µg/L total lead).^22^ The metal mixture was comprised of all four metals, each at the same concentration as individual metal exposures. F1 offspring were weaned on postnatal day 21 and subsequently provided untreated water for two weeks until tissue collection.

### Tissue Collection and RNA Extraction

At the end of the exposure period, offspring were euthanized and hearts were excised and weighed. Hearts were rinsed briefly with phosphate-buffered saline (PBS) to remove residual blood, halved, and snap frozen in liquid nitrogen for subsequent RNA extraction. Total RNA was extracted from one half of each heart using the AllPrep DNA/RNA Mini Kit (Qiagen, Cat#80204, Germantown, Maryland) according to manufacturer’s instructions. RNA concentration was measured using spectrophotometry (NanoDrop 100, ThermoFisher). Bulk RNA was then sent to Novogene Corporation, Inc. (Novogene, Beaverton, OR) for sample quality control, library preparation, and sequencing. A total of six samples per sex and exposure group were sent for sequencing.

### Novogene info: Library Prep and Sequencing

All sample quality check, library construction, and sequencing was completed by Novogene using standard protocols. Briefly, RNA quality was determined using a combination of 1% agarose gel electrophoresis, NanoDrop spectrophotometry, and RNA Integrating Number confirmation with Agilent2100 measurements. Following quality checks, samples meeting quality criteria were used for library construction. Sequencing libraries were prepared using polyA enrichment and were subsequently sequenced using the 150-bp paired end sequencing (PE150) strategy offered through Novogene. Once sequencing was completed, the resulting gene count files were used for all downstream bioinformatics analysis.

### Differential Expression and Functional Enrichment

First, outlier samples were identified before examining treatment effects using principal component analysis (PCA) and calculating the Mahalanobis distance; these outliers (HT_Pb_M6, HT_Pb_F5, HT_Cd_M6, HT_Cr_M5) were then removed from downstream analysis (data included in **Supplemental Figure 1**). Differential expression analysis was performed using a combination of *DESeq2*^23^ (through the *tidybulk* package^24^) and bootstrapping with replacement (number of iterations = 100). Two statistical models were employed for the results of this study:

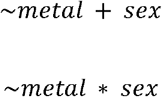

Both statistical models used the untreated control as the reference level for metal and males as the reference level for sex. In both models, bootstrapping results were filtered by two metrics: (1) how frequently a gene was differentially expressed (median nominal *P-value* < *0.05* and median |Log_2_-fold change| >*0.25*) and (2) the sign change (defined as the amount of times the direction that differentially expressed gene (DEG) flipped to the opposite direction < *0.25*). In short, we considered genes that were significant at least 30% of the time and switched between up- and downregulation less than 25% of the time, as these were considered the most stable across bootstrapping iterations. Significant DEGs were then analyzed for functional enrichment through *enrichGO* (part of the *clusterProfiler* package^25^) to identify significant gene ontology (GO) terms. These individual terms were then processed to identify higher level parent terms using the *rrvgo* package^26^ for each metal treatment group and each sex (for interaction model only). We assessed the GO term overlap using an enrichment score:

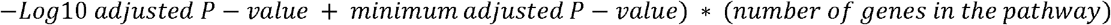

When GO terms were collapsed into parent GO terms, the median enrichment score of the collapsed GO terms was calculated and reported.

### Deconvolution and DGCA

To perform deconvolution, we pseudobulked^24^ a publicly-available mouse heart single-cell reference dataset from CellxGene^27^ using the *Seurat*^28^ package. This reference contained single-cell data from control mice and mice subjected to heart injury. Only the control (sham) mice were included for deconvolution. Cell type proportion changes were assessed using linear modeling and considered significant if the *P-value* was below *0.1*:

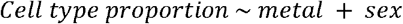

After performing deconvolution, we used differential gene correlation analysis (*DGCA* package^29^) to understand if the relationship between the cell profiles changed after metal treatment. We compared cell type relationships between each metal compared to the untreated control. Cellular relationships were considered perturbed if the empirical *P-value* was below *0.1*.

### GSVA Analysis

Gene set variation analysis (*GSVA*) was performed using the *msigdbr*^30^ mouse category C5 gene ontology annotated pathways for postnatal heart growth and maturation. Gene sets were constructed for key heart development pathways and *GSVA* was performed using the *GSVA* package^31^ for each metal treatment group. To explore relationships between pathway activity and cellular composition, linear regression was performed between the GSVA score and cell type proportion change from the deconvolution results for each metal. Both GSVA and linear regression were performed with the following model:

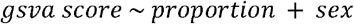

Significant results were considered with a nominal *P-value* < *0.05*.

## Results

### Heavy metal exposure alters gene expression in a metal-specific manner

To determine how perinatal exposure to heavy metals affects the cardiac transcriptome with our covariate model, differentially expressed genes (DEGs) were visualized using the *EnhancedVolcano* package,^32^ where red dots indicate significantly upregulated DEGs, blue dots indicate significantly downregulated DEGs, and black dots indicate non-significant DEGs (**Figure 1A**). Noteworthy genes that changed with different individual metal exposures include *Arntl* or *Bmal1* (Brain and Muscle ARNT-Like 1, arsenic and chromium)*, Spon2* (Spondin-2, cadmium and chromium), and in the mixture group, *Nppb* (B-type natriuretic peptide). In order to determine how each metal treatment overlapped with the transcriptome changes associated with the mixture group, circos plots were generated (using the *circlize* package^33^) to show the percent of each metal’s total significant DEGs shared with the mixture, indicated by the darker coloring of the ramps (**Figure 1B**). Lead shared the greatest percentage of its significant DEGs with the mixture (∼38.7%), while cadmium shared the smallest percentage (∼27.2%) (**Figure 1B**). In order to determine functional enrichment of our significant DEGs, we utilized gene ontology biological process (GO BP) and mapped to parent terms (**Figure 1C**). Median enrichment score was calculated by taking -Log_10_(*P-value* + 1) multiplied by the count (the number of DEGs in that pathway). All GO terms for each condition, as well as the parent terms they mapped to are included in **Supplemental Table 1**. Parent GO terms showed metal-specific directional enrichment in terms relating to nuclear division, immune signaling, and circadian rhythm (**Figure 1C**). Arsenic was associated with downregulation of pathways involved in cellular division and circadian rhythm of gene expression and upregulated regulation of muscle contraction, while cadmium enriched pathways in cellular response to fibroblast growth factor stimulus, neurodevelopment pathways, and inorganic anion transport (**Figure 1C**). Chromium led to a downregulation of pathways related to neurodevelopment, circadian rhythm, and double-stranded break repair (**Figure 1C**). In contrast, lead upregulated pathways associated with cellular division and membrane-less organelle assembly (**Figure 1C**). Interestingly, exposure to the metal mixture only mapped to circadian rhythm of gene expression (**Figure 1C**). Collectively, these results suggest that transcriptome-level changes are metal-specific and heavy metal exposure alters pathways associated with circadian rhythm, heart function, and DNA damage repair.

**Figure 1:**
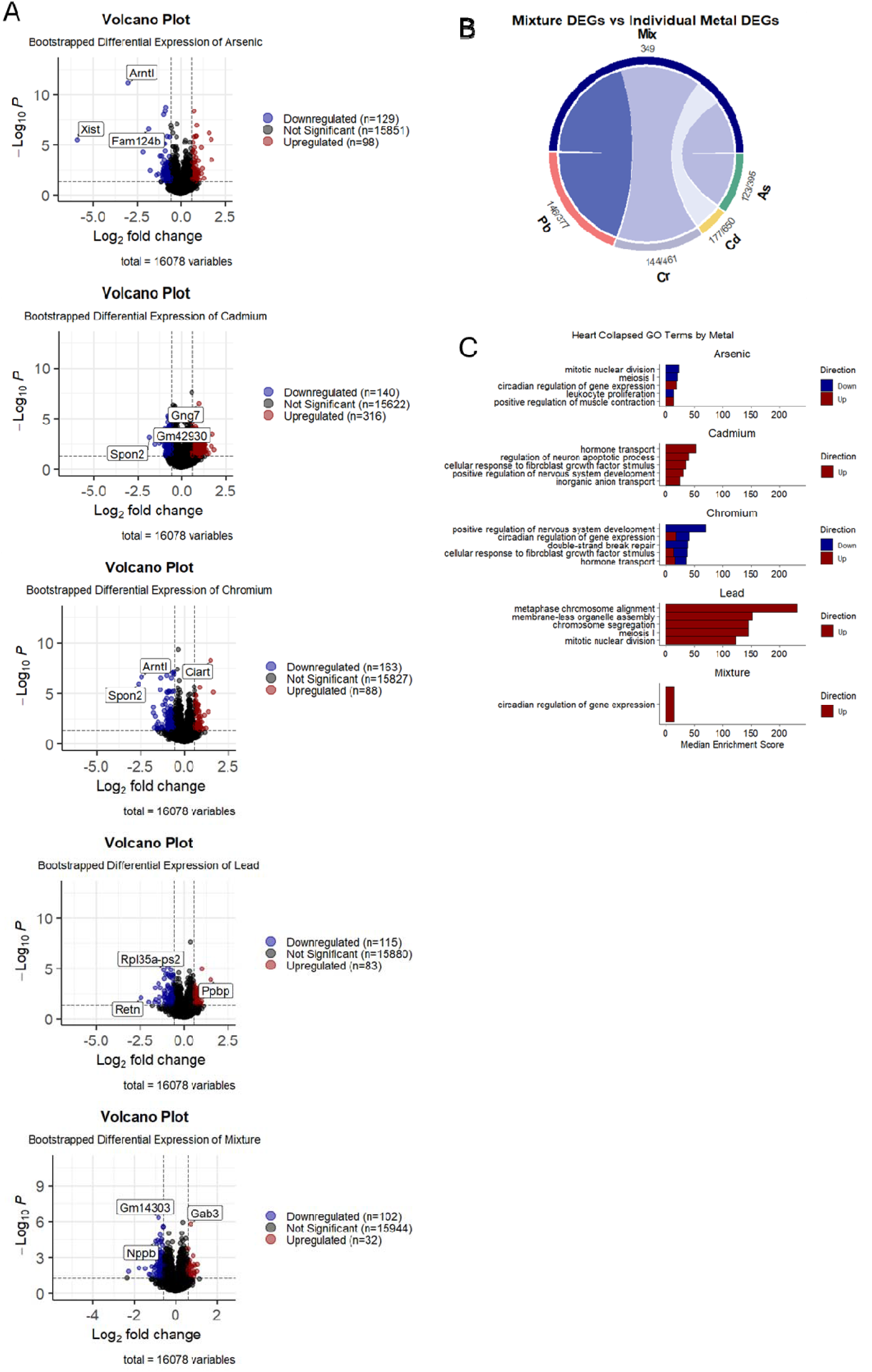
Summary of differential expression and GO enrichment analysis. A) Volcano plots showing total DEGs between for arsenic, cadmium, chromium, lead, and mixture. B) Circos plots indicating percentage of DEGs shared between each individual metal treatment and the mixture. C) Top five GO enrichment terms for each metal treatment group ordered by median enrichment score (calculated as -log_10_(*P_adj_* + min(*P_adj_*) * (number of genes). Red indicates upregulation of that pathway and blue indicates downregulation of that pathway term.

### Heavy metal exposure alters normal cardiac cell composition and triggers NK cell activation

To determine potential cardiac cell proportion perturbations associated with perinatal heavy metal exposure, we used deconvolution analysis (**Figure 2A)**, where red indicates cell types increased with specific exposures and blue indicates cell types that are decreased. Overall, arsenic, cadmium, chromium, and mixture exposure increased fibroblast proportions in both sexes, and all treatment groups except chromium, led to a decrease in cardiac muscle cell proportions compared to the untreated group (**Figure 2A**). To determine concordance in cell type proportion directional changes consistent with the mixture group, we quantified the number of cell type proportion changes in the same direction of the mixture as the individual metals out of the total number of cell types (n=7) (**Figure 2B**). Cadmium was most concordant with the mixture in cell type proportion changes, with 6 out of the 7 cell types changing in the same direction, while arsenic had the lowest concordance with the mixture. Next, we performed DGCA to understand how cellular relationships changed after metal exposure, where a relationship was deemed perturbed if the empirical *P-value* was below 0.1. Raw DGCA results can be found in **Supplemental Figure 2**. Dendritic cells had the greatest number of significantly altered relationships, followed by cardiac muscle cells, natural killer (NK) cells, and endothelial cells. Given that relationships with immune cell profiles were consistently perturbed, we looked at the gene expression of targeted genes for general surface markers (*Cd11c/Itgax* and *Cd56/Ncam1* for dendritic cells and NK cells, respectively) and for activation markers of these cell types (*Cd40/Tnfrsf5* & *Cd86* for dendritic cells, *Cd69* & *Cd25/Il2ra* for NK cells) (**Figure 2D-I**). No significant changes were observed in surface marker or activation marker gene expression in dendritic cells. However, there was a significant increase in the general surface markers for NK cells with cadmium treatment (**Figure 2G**) and a significant increase in *Cd25* expression with arsenic and mixture exposure, with trending increases in this marker with cadmium and chromium (**Figure 2I**). Together, these findings suggest that perinatal heavy metal exposure may alter the normal cellular composition of the postnatal heart and activate NK cells, an important component of the cardiac immune system.

**Figure 2:**
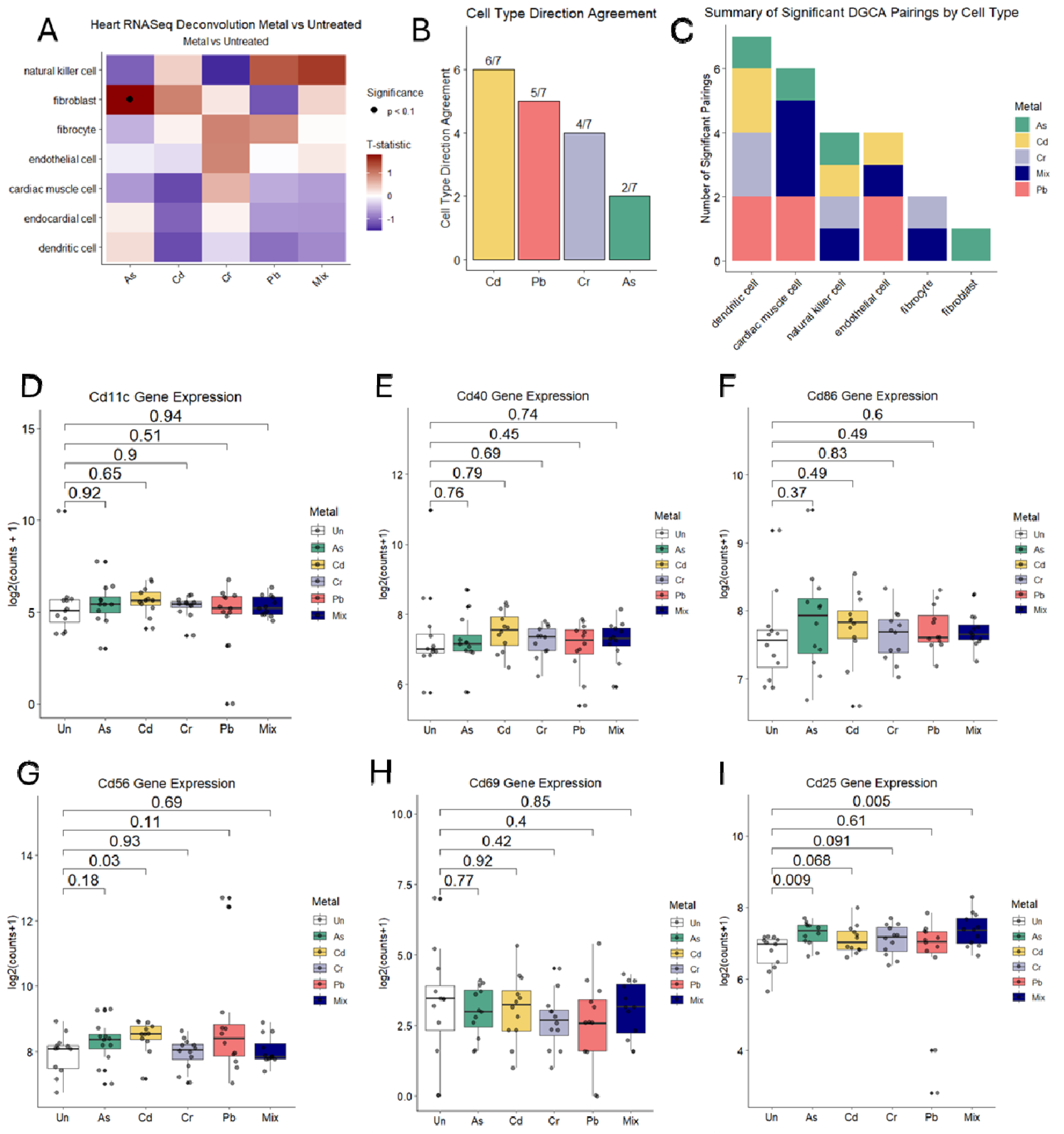
Deconvolution and DGCA results for cardiac cell type changes. A) Summary of deconvolution proportion changes for metal treatment groups. B) Cell type proportion change agreement of each individual metal with the mixture. C) Summary of number of significant DGCA pairings for each cell type across treatment groups. D) Dendritic cell surface marker (*Cd11c*) gene expression for each treatment group, E & F) Dendritic cell activation marker (*Cd40* & *Cd86*) gene expression for each treatment group. G) NK cell surface marker (*Cd56*) gene expression for each treatment group. H & I) NK cell activation marker (*Cd69* & *Cd25*) gene expression for each treatment group.

### Perinatal metal exposure alters pathways related to postnatal heart growth and metabolic maturation

Following the cell type proportion changes identified in **Figure 2**, we targeted postnatal heart maturation pathways with individual metal and metal mixture exposure using GSVA (**Figure 3A**), where red indicates an increase in that pathway, blue indicates a decrease in that pathway, and dots indicate statistically significant changes (*P-value* < 0.05). A complete list of GO BP terms and their targeted pathways is included in **Supplemental Table 2**. Overall, metal mixture exposure led to a general upregulation of postnatal heart maturation pathways, including growth and developmental signaling, vascular development, and calcium handling and electrophysiology. Additionally, we saw a slight downregulation of metabolic maturation pathways with cadmium, lead, and the mixture. These results highlight the potential for an energetic mismatch in maturing postnatal hearts. Next, we wanted to determine whether the cell type proportion changes in our deconvolution results correlated with our GSVA postnatal heart maturation pathways, so we performed a linear regression between the deconvolution cell type proportion and the GSVA pathway score for each metal compared to the untreated control (**Figure 3B**). In general, cell type proportion changes were inversely associated with GSVA pathways. Two exceptions to this trend were changes to fibroblasts and fibrocytes, which were generally positively associated with GSVA pathways. Taken together, these results suggest that heavy metal exposure upregulates postnatal heart growth and maturation, but favors downregulation of metabolic maturation pathways.

**Figure 3:**
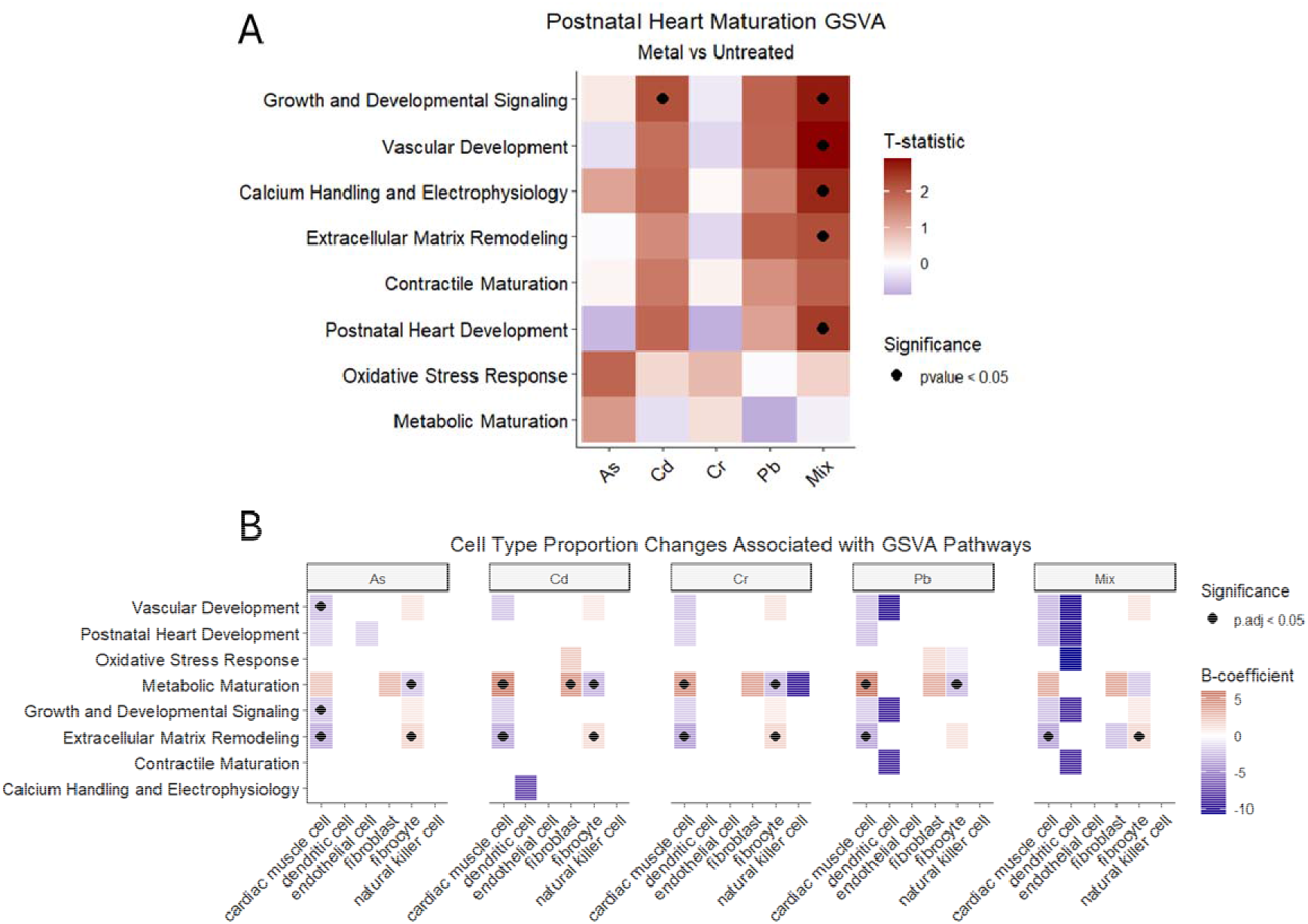
Postnatal heart maturation alterations due to metal exposure. A) Postnatal heart maturation GSVA results for individual treatment groups, where red indicates an increase in that pathway and blue indicates a decrease in that pathway. B) Linear regression results between cell type proportion changes from deconvolution and GSVA pathway scores faceted by metal treatment group.

### Metal exposure induces sex-specific genetic changes and alters immune-related pathways in males

Finally, we repeated our differential expression analysis from **Figure 1** modeling sex as an interaction term by each treatment to identify sex-dependent transcriptional responses. We again employed a combination of bootstrapping with replacement and differential expression with *DESeq2*^23^ with this model and separated by female- and male-specific differentially expressed genes (DEGs). DEGs were visualized in volcano plots for both female (left) and male (right) DEGs (**Figure 4A**). Notably, both arsenic and chromium downregulated *Bmal1/Arntl* (Brain and Muscle ARNT-Like 1) and cadmium and chromium upregulated *Dbp* (D-box binding PAR bZIP transcription factor). In males, there was less overlap between treatments, but chromium and lead downregulated *Nr4a3* (nuclear receptor subfamily 4 group A member 3). Chromium shared the greatest percentage of its significant DEGs with the mixture in female-specific DEGs (∼17.1%) and arsenic shared the greatest percentage of its significant DEGs with the mixture in male-specific DEGs (∼8.3%) (**Figure 4B**). However, there were far fewer male-specific DEGs than female-specific DEGs, so this overlap is minimal. We again performed functional enrichment of our significant DEGs with GO BP mapping to identify parent GO terms involved in metal treatment exposures for females (left) and males (right) (**Figure 4C**). Female-specific pathways had more metal-specific enrichment. Arsenic upregulated pathways associated with cellular energy production (pyridine-containing compound metabolic process), GTPase-mediated signaling, and connective tissue development. Cadmium upregulated pathways relating to rhythmic processes, positive regulation of muscle contraction, and blood vessel diameter regulation (**Figure 4C**). Chromium upregulated pathways involved in regulation of inflammatory response, lymphocyte differentiation, and negative regulation of lymphocyte apoptosis (**Figure 4C**). Lead upregulated pathways involved in T cell proliferation, leukocyte adhesion and chemotaxis, and lymphocyte differentiation (**Figure 4C**). Metal mixture exposure upregulated pathways involved in nuclear division, rhythmic processes, and inflammatory response mediation. There was more of an immune focus in male-specific DEGs (**Figure 4C**). Arsenic, lead, and metal mixture exposures downregulated terms related to antigen processing and presentation (**Figure 4C**). Arsenic, chromium, and mixture exposures downregulated terms relating to leukocyte activation, and chromium, lead, and mixture exposures downregulated terms relating to leukocyte adhesion (**Figure 4C**). Together, these results indicate that heavy metal exposure has sex-specific transcriptome-level effects and favors the downregulation of immune-related terms in males, but not females.

**Figure 4:**
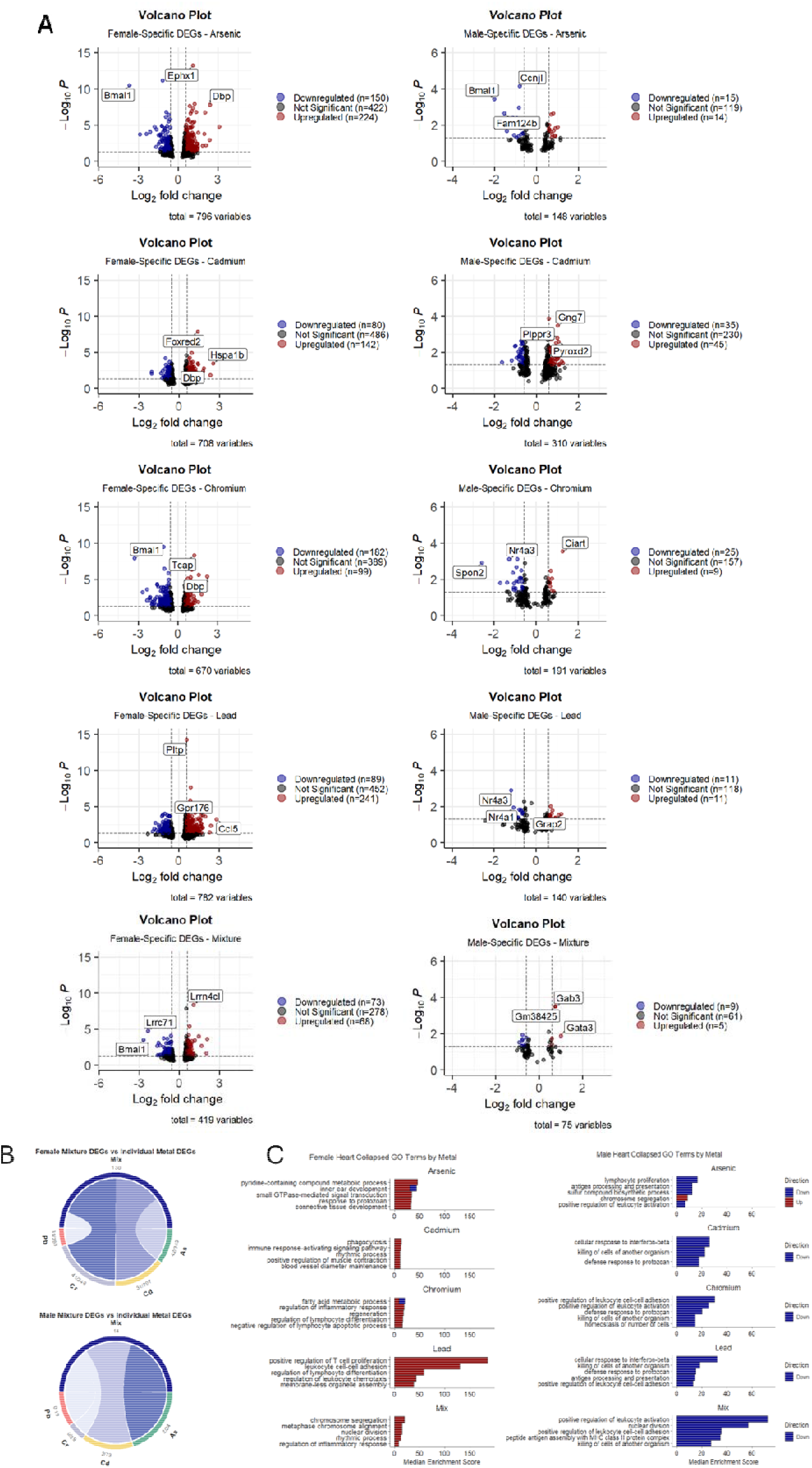
Differential gene expression and GO enrichment results for interaction model. A) Volcano plots showing total DEGs between for arsenic, cadmium, chromium, lead, and mixture in both females (left) and males (right). B) Circos plot indicating percentage of DEGs shared between each individual metal treatment groups and the mixture for females (top) and males (bottom). C) Top five GO enrichment terms for each metal treatment group ordered by median enrichment score (calculated as -log_10_(*P_adj_*+ min(*P_adj_*) * (number of genes) for females (left) and males (right). Red indicates upregulation of that pathway and blue indicates downregulation of that pathway term.

## Discussion

Herein, we characterized the effects of perinatal heavy metal and metal mixture exposure on the offspring cardiac transcriptome for the first time. These metals are thought to impair the formation and maturation of cardiac muscle cells,^34–36^ particularly during early gestation when the heart is most vulnerable.^37^ A common mechanism of toxicity is the generation of reactive oxygen species, leading to DNA damage, protein oxidation, and mitochondrial dysfunction.^38^ In this study, we found that metal exposure altered gene expression related to heart rate regulation and cell division, changed cardiac cellular composition, caused an energetic-maturation mismatch, and increased resilience to metal-induced damage in females.

### Heavy metal exposure alters gene expression related to heart rate regulation and cell division

Exposure to individual metals and the metal mixture altered the cardiac transcriptional landscape of offspring, particularly genes related to heart function and nuclear division. Perinatal metal exposure downregulated *Arntl* (*Bmal1*, arsenic and chromium), a circadian regulator,^39^ *Spon2* (cadmium and chromium), a cardioprotective immune signaling gene,^40,41^ and *Nppb* (mixture only), which regulates blood pressure and protects against cardiac stress^42^ (**Figure 1A**). These findings suggest heavy metals may promote cardiac dysfunction by suppressing cardioprotective pathways. Although reduced *Arntl/Bmal1* expression has been reported following cadmium^43^ and lead^44,45^ exposure, arsenic has been shown to increase *Arntl/Bmal1* expression in urothelial carcinoma cells,^46^ underscoring tissue- and context-dependent effects. Arsenic has been shown to decrease *Spon2* expression in macrophages,^47^ but comparable studies for other metals are lacking. In contrast, arsenic,^48,49^ cadmium,^50–52^ chromium (total),^53^ and lead^52,54^ have all been associated with increased *Nppb* expression, although the effects of metal mixtures and gestational exposure remain unstudied. Since circadian rhythm regulates heart rate and conduction,^55^ disruption of these pathways may have lasting effects on cardiac function. Overall, heavy metal exposure predominantly downregulated gene expression, except for cadmium, which induced more upregulated than downregulated genes (**Figure 1A**). However, whether heavy metal exposure preferentially downregulates or upregulates gene expression remains inconsistent across studies. For example, arsenic has been reported to favor downregulation in bladder epithelial cells,^56^ but upregulation in HeLa cells,^57^ highlighting the context dependent nature of its transcriptional effects. Moreover, we found that lead shared the greatest proportion of DEGs with the mixture (**Figure 1B**), suggesting similar transcriptional responses, although this was not reflected in the GO enrichment results (**Figure 1C**). This discrepancy could reflect differences in the concordance of shared genes vs. enriched pathways. Heavy metal exposure also altered genes involved in nuclear division (**Figure 1C**), consistent with evidence that metal-induced DNA damage impairs cellular division.^58^ Arsenic and chromium reduced pathways related to nuclear division and double-stranded break repair (**Figure 1C**), suggesting they not only disrupt proliferation but also compromise DNA repair, which is consistent with prior work.^59^ Collectively, these findings indicate that perinatal heavy metal exposure disrupts key pathways regulating circadian rhythm, cell division, and DNA repair, with potential consequences for cardiac structure and function in offspring.

### Metal exposure disrupts normal cardiac cellular composition

Our deconvolution analysis showed that heavy metal exposure generally decreased cardiomyocyte and dendritic cell proportions, while increasing fibroblast and fibrocyte populations (**Figure 2A**). These changes were metal-specific, with cadmium most closely resembling the mixture (**Figure 2B**). Consistent with previous studies, arsenic,^60^ cadmium,^61^ chromium,^62^ and lead^63^ have all been linked to increased cardiomyocyte apoptosis through oxidative stress^61,62^ or reduced antioxidant capacity,^60,63^ which may impair cardiac function. GO enrichment showed increased expression of genes involved in muscle contraction and ion transport, possibly reflecting a compensatory response to cardiomyocyte loss. Heavy metals, including arsenic and cadmium, have also been linked to cardiac fibrosis in adults.^5,64,65^ Although lead has been associated with cardiac fibrosis, our deconvolution analysis instead showed a slight decrease in fibroblasts but a slight increase in fibrocytes following lead exposure (**Figure 2A**). These findings differ from previous work showing that early-life lead exposure increased cardiac fibrosis in adulthood,^66^ possibly reflecting differences in dose (50mg/kg/day), exposure method (gavage), developmental time, or species (rats). Chromium cardiotoxicity is less well-characterized but has been linked to increased heart rate in zebrafish embryos,^67^ as well as oxidative stress,^68^ a known driver of cardiac fibrosis.^69^ We also observed an overall decrease in cardiomyocytes following heavy metal exposure, with the exception of chromium (**Figure 2A**). An increase in fibrosis-associated cells, together with cardiomyocyte loss, could increase cardiac stiffness and impair function. We also observed altered relationships between dendritic and NK cells (**Figure 2C**). While dendritic cell marker expression was unchanged, cadmium increased NK cell surface marker expression, and arsenic and the metal mixture increased expression of activation marker *Cd25* (**Figures 2D-I**). NK cells help prevent excess collagen deposition and cardiac fibrosis,^70^ so their increased expression may represent a response to cardiomyocyte loss and increased fibrotic signaling following metal exposure. This aligns with studies linking heavy metal exposure to cardiomyocyte apoptosis, endothelial cell dysfunction, and immune activation.^71^ Collectively, perinatal heavy metal exposure alters cardiac cell composition by reducing cardiomyocytes, and increasing fibroblast and fibrocyte populations, potentially impairing cardiac function and increasing stiffness.

### Metal mixture exposure introduces a potential transcriptional energetic-maturation mismatch

To determine whether heavy metal exposure disrupts cardiac maturation, we examined pathways involved in calcium signaling,^72,73^ metabolic maturation,^74,75^ and vascular development.^76^ Previous studies show these pathways peak at four weeks post-natal,^77^ matching the age of mice in this study. Heavy metal exposure increased growth, vascular development, and calcium handling pathways, while reducing metabolic maturation pathways. The metal mixture produced an overall increase in most of our heart maturation pathways (**Figure 3A**), suggesting a potential energetic-maturation mismatch. As such, individual metals at regulatory levels may have limited effects on postnatal cardiac maturation, but combined exposure as a mixture significantly alters cardiac maturation pathways. Increased developmental signaling alongside reduced metabolic maturation suggests premature maturation and an energetic-maturation mismatch. This may result from impaired metabolic transitions, as metals like arsenic, can disrupt fatty acid oxidation,^78^ a key energy source for mature cardiomyocytes.^74,75^ The mixture effects appear from combined metal exposure rather than a single driving metal. Cadmium and lead can disrupt calcium handling,^79–81^ a key component of cardiac contraction, consistent with our findings of reduced cardiomyocytes and increased fibrotic cell populations. Correlation of cell composition and pathway changes showed that cardiomyocyte proportions were inversely associated with growth, vascular development, and extracellular matrix remodeling, but positively associated with metabolic maturation across treatment groups (**Figure 3B**). In contrast, fibrocytes were positively associated with extracellular matrix remodeling (**Figure 3B**), consistent with their role in maintaining tissue structure.^82^ Together, these findings suggest that perinatal heavy metal exposure alters cardiac cell composition and disrupts maturation pathways important for long-term cardiovascular health.

### Heavy metals alter gene expression in a sex-specific manner and reveal that females are resilient to metal-induced damage

Using an interaction model, we found that heavy metal exposure preferentially upregulated female-specific genes and downregulated male-specific genes (**Figure 4A**). In females, heavy metal exposure consistently decreased *Bmal1* (or *Arntl*) and increased *Dbp*, key circadian regulators.^83,84^ Males exhibited fewer DEGs, although chromium and lead both reduced *Nr4a3*, a gene linked to cardiovascular disease.^85^ Moreover, chromium shared the greatest overlap with the mixture in females, whereas arsenic shared the greatest overlap in males (**Figure 4B**). Male-enriched pathways were primarily related to immune function, including lymphocyte proliferation, leukocyte adhesion, and leukocyte activation (**Figure 4C**). In contrast, female-specific genes were enriched for pathways involved in heart function and repair, including muscle contraction, rhythmic process, and regeneration, suggesting a compensatory response to heavy metal-induced damage. A full list of female- and male-specific GO terms can be found in **Supplemental Table 3**. These findings suggest that females are more resilient to metal-induced cardiac damage than males. Indeed, we previously found that arsenic and cadmium impaired cardiac structure and function in adult male mice but not females.^7,8^ Males have also been shown by others to be more sensitive to metal-induced injury compared to females.^86^ Female-specific pathways also showed increased immune response terms, contrasting with the immune-related changes observed in males. Previous studies have reported both immune activation and immunosuppression following heavy metal exposure, indicating context-dependent immune effects.^87–89^ As such, our current findings align with prior work showing greater susceptibility to metal-induced cardiac injury in males than females. Overall, these results demonstrate sex-specific cardiac transcriptome responses, with females exhibiting greater adaptive capacity to metal-induced cardiac damage.

### Limitations

Although this is among the first studies to examine perinatal heavy metal mixture-associated changes in the cardiac transcriptome, there are multiple limitations that require acknowledgement. Firstly, we assessed only a single postnatal timepoint after exposure ended, so acute effects may have been missed despite the persistent transcriptomic changes observed. Future studies should examine additional prenatal and postnatal timepoints to better define how metal exposure disrupts gene expression during critical stages of cardiac development. This study also used bulk-RNA sequencing, so future studies should validate our deconvolution findings with single-cell RNA sequencing. Protein-level analyses and cardiac functional measurements are also needed to determine the impact of the observed transcriptomic changes. Finally, evaluating multiple exposure doses will be important to define dose-response relationships for metal exposures in the developing heart.

## Conclusions

Overall, perinatal exposure to heavy metals at regulatory limits induced metal- and sex-specific changes in the postnatal cardiac transcriptome. Heavy metal exposure altered pathways related to circadian rhythm, nuclear division, immune function, cardiac cell composition, and postnatal heart maturation. We also identified increased developmental signaling without corresponding metabolic maturation, suggesting a potential energetic-maturation mismatch in the postnatal heart. Females preferentially upregulated genes in cardiac structure and function, whereas males downregulated immune-related genes, suggesting greater female resilience to heavy metal-induced damage. The metal mixture also produced a distinct transcriptomic profile, indicating that combined exposure may be more harmful than individual metals, thus underscoring the need for mixture-focused toxicological studies. Together, these findings support early development as a critical window of susceptibility, with effects that persist beyond the exposure period.

## Acknowledgements

We acknowledge the authors of the original NEMMEX manuscript^19^ for leading the *in vivo* experiments that yielded biospecimens for this follow-up study.

## Declaration of Generative AI and AI-assisted Technologies in the writing process

Generative AI and AI-assisted technologies were not used in the preparation of this work, and the authors take full responsibility for the content of the final manuscript.

## CRediT authorship contribution statement

**Morgan Steiner:** Conceptualization; Investigation; Data curation; Formal analysis; Visualization; Writing – original draft; Writing – review & editing. **Jason Laird:** Formal analysis; Writing – review & editing. **Sylvia S. Sanchez:** Investigation; Writing – review & editing. **Susmitha Pagadala:** Investigation. **Shyam Biswal:** Funding acquisition; Writing – review & editing. **Fenna C. M. Sillé:** Conceptualization; Investigation; Resources; Funding acquisition; Supervision; Project administration; Writing – original draft; Writing – review & editing. **Mark J. Kohr:** Conceptualization; Investigation; Supervision; Writing – original draft; Writing – review & editing.

## Supplementary Figure Legends

**Supplemental Figure 1:**
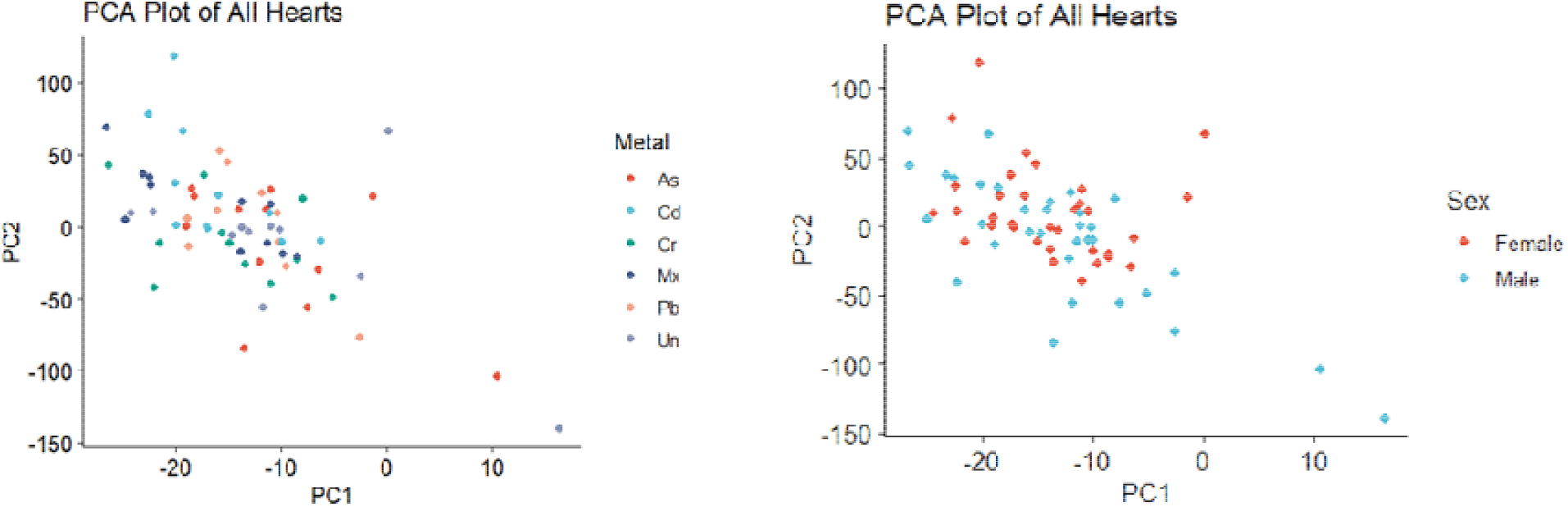
PCA plots of heart samples by metal and sex. PCA plots were generated from all heart samples and colored by metal (left) or sex (right). These PCA plots are after the removal of four outliers total across all groups.

**Supplemental Figure 2:**
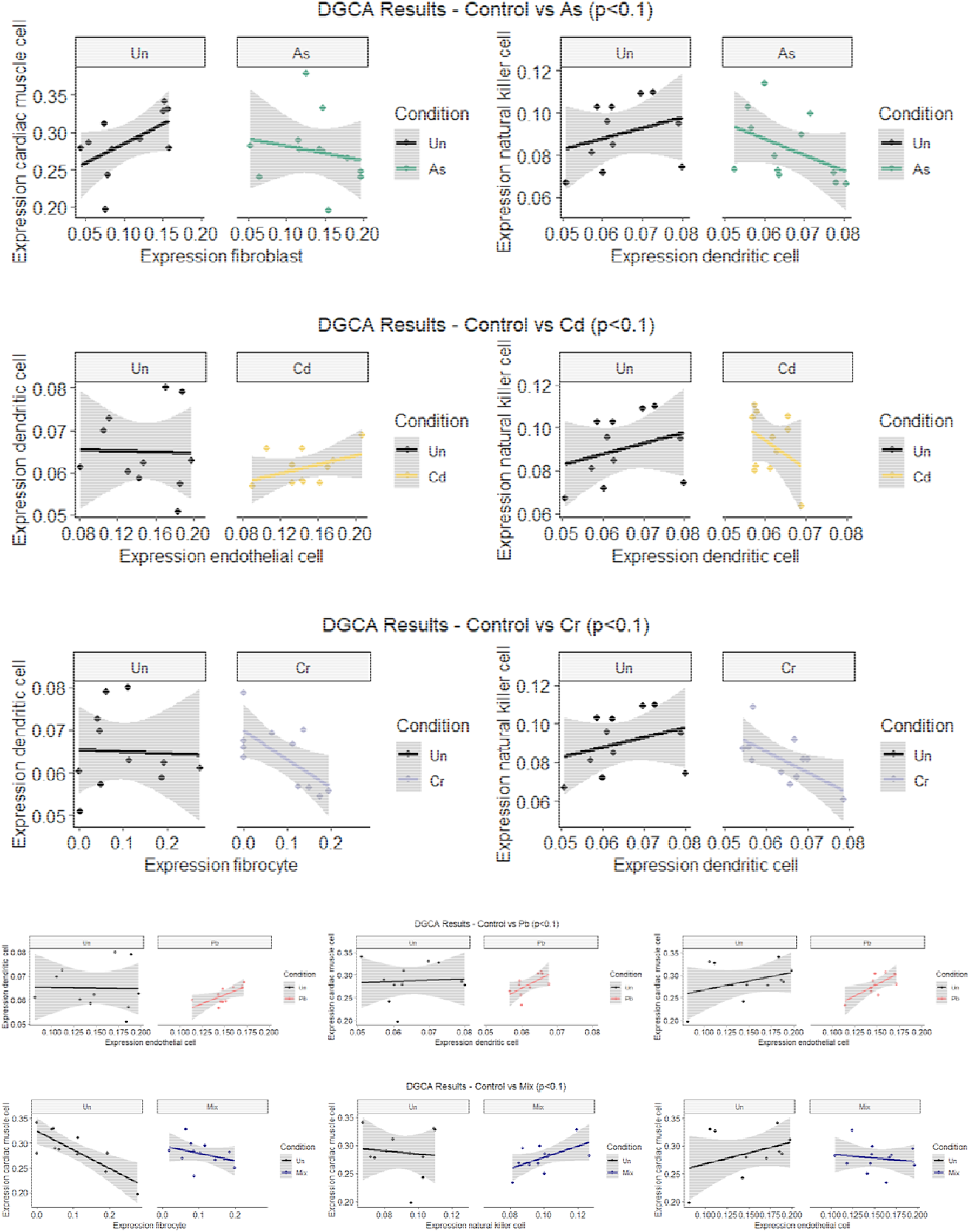
Significant DGCA pairings for all metals. DGCA pairings were filtered with a significance cutoff of p<0.1.

## Supplementary Table Legends

**Supplemental Table 1:** Complete list of GO BP terms and mapped parent terms by metal (see Excel sheet).

**Supplemental Table 2:** GSVA pathway targeted GO:BP terms for postnatal heart development and maturation (see Excel sheet).

**Supplemental Table 3:** Female- and male-specific GO term enrichment results (see Excel sheet).

## References

1. Briffa J, Sinagra E, Blundell R. Heavy metal pollution in the environment and their toxicological effects on humans. Heliyon. 2020;6(9):e04691. doi:10.1016/j.heliyon.2020.e04691

2. Vareda JP, Valente AJM, Durães L. Assessment of heavy metal pollution from anthropogenic activities and remediation strategies: A review. J Environ Manage. 2019;246:101–118. doi:10.1016/j.jenvman.2019.05.126

3. Masindi V, Muedi KL. Environmental contamination by heavy metals. Heavy Met. 2018;10(4):115–133.

4. Yang A, Lo K, Zheng T, et al. Environmental heavy metals and cardiovascular diseases: Status and future direction. Chronic Dis Transl Med. 2020;6(4):251–259. doi:10.1016/j.cdtm.2020.02.005

5. Hocher B. Metal exposure in heart disease. Clin Chim Acta. 2026;589:121011. doi:10.1016/j.cca.2026.121011

6. Veenema R, Casin KM, Sinha P, et al. Inorganic arsenic exposure induces sex-disparate effects and exacerbates ischemia-reperfusion injury in the female heart. Am J Physiol-Heart Circ Physiol. 2019;316(5):H1053–H1064. doi:10.1152/ajpheart.00364.2018

7. Kabir R, Sinha P, Mishra S, et al. Inorganic arsenic induces sex-dependent pathological hypertrophy in the heart. Am J Physiol-Heart Circ Physiol. 2021;320(4):H1321–H1336. doi:10.1152/ajpheart.00435.2020

8. Fitch ML, Kabir R, Ebenebe OV, et al. Cadmium exposure induces a sex-dependent decline in left ventricular cardiac function. Life Sci. 2023;324:121712. doi:10.1016/j.lfs.2023.121712

9. Taube N, Kabir R, Ebenebe OV, et al. Prenatal arsenite exposure alters maternal cardiac remodeling during late pregnancy. Toxicol Appl Pharmacol. 2024;483:116833. doi:10.1016/j.taap.2024.116833

10. Taube N, Steiner M, Ebenebe-Kasonde OV, et al. Gestational arsenite exposure alters maternal postpartum heart size and induces Ca^2+^ -handling dysregulation in cardiomyocytes. Am J Physiol-Heart Circ Physiol. 2025;328(3):H460–H471. doi:10.1152/ajpheart.00266.2024

11. Suhl J, Conway KM, Rhoads A, et al. Prepregnancy exposure to dietary arsenic and congenital heart defects. Birth Defects Res. 2023;115(1):79–87. doi:10.1002/bdr2.2110

12. Jin X, Tian X, Liu Z, et al. Maternal exposure to arsenic and cadmium and the risk of congenital heart defects in offspring. Reprod Toxicol. 2016;59:109–116. doi:10.1016/j.reprotox.2015.12.007

13. Liu Z, Yu Y, Li X, et al. Maternal lead exposure and risk of congenital heart defects occurrence in offspring. Reprod Toxicol. 2015;51:1–6. doi:10.1016/j.reprotox.2014.11.002

14. Chen Y, Wu F, Liu X, et al. Early life and adolescent arsenic exposure from drinking water and blood pressure in adolescence. Environ Res. 2019;178:108681. doi:10.1016/j.envres.2019.108681

15. Farzan SF, Howe CG, Chen Y, et al. Prenatal lead exposure and elevated blood pressure in children. Environ Int. 2018;121(Pt 2):1289–1296. doi:10.1016/j.envint.2018.10.049

16. Liu Y, Yu L, Zhu M, et al. Associations of exposure to multiple metals with blood pressure and hypertension: A cross-sectional study in Chinese preschool children. Chemosphere. 2022;307:135985. doi:10.1016/j.chemosphere.2022.135985

17. Son J, Morris JS, Park K. Toenail Chromium Concentration and Metabolic Syndrome among Korean Adults. Int J Environ Res Public Health. 2018;15(4):682. doi:10.3390/ijerph15040682

18. Sazakli E, Villanueva C, Kogevinas M, Maltezis K, Mouzaki A, Leotsinidis M. Chromium in Drinking Water: Association with Biomarkers of Exposure and Effect. Int J Environ Res Public Health. 2014;11(10):10125–10145. doi:10.3390/ijerph111010125

19. Chandra N, Karimi B, Bhobe A, et al. Perinatal exposure to metal mixtures disrupts neuronal function and behavior. Environ Res. 2026;294:123803. doi:10.1016/j.envres.2026.123803

20. Percie Du Sert N, Hurst V, Ahluwalia A, et al. The ARRIVE guidelines 2.0: Updated guidelines for reporting animal research. Br J Pharmacol. 2020;177(16):3617–3624. doi:10.1111/bph.15193

21. EPA US. National Primary Drinking Water Regulations; Synthetic Organic Chemicals and Inorganic Chemicals; Monitoring for Unregulated Contaminants; National Primary Drinking Water Regulations Implementation; National Secondary Drinking Water Regulations. Final Rule, January 31, 1991. 1991.

22. U.S. EPA. National Primary Drinking Water Regulations; Synthetic Organic Chemicals and Inorganic Chemicals; Monitoring for Unregulated Contaminants; National Primary Drinking Water Regulations Implementation; National Secondary Drinking Water Regulations. Final Rule, January 31, 1991. *FedReg*. 1991;Federal Register 56(3526-3597):31776.

23. Love MI, Huber W, Anders S. Moderated estimation of fold change and dispersion for RNA-seq data with DESeq2. Genome Biol. 2014;15(12):550. doi:10.1186/s13059-014-0550-8

24. Mangiola S, Molania R, Dong R, Doyle MA, Papenfuss AT. tidybulk: an R tidy framework for modular transcriptomic data analysis. Genome Biol. 2021;22(1):42. doi:10.1186/s13059-020-02233-7

25. Xu S, Hu E, Cai Y, et al. Using clusterProfiler to characterize multiomics data. Nat Protoc. 2024;19(11):3292–3320. doi:10.1038/s41596-024-01020-z

26. Sayols S. rrvgo: a Bioconductor package for interpreting lists of Gene Ontology terms. MicroPublication Biol. 2023;2023. doi:10.17912/micropub.biology.000811

27. Cortada E, Yao J, Xia Y, et al. Cross-species single-cell RNA-seq analysis reveals disparate and conserved cardiac and extracardiac inflammatory responses upon heart injury. Commun Biol. 2024;7(1):1611. doi:10.1038/s42003-024-07315-x

28. Hao Y, Stuart T, Kowalski MH, et al. Dictionary learning for integrative, multimodal and scalable single-cell analysis. Nat Biotechnol. 2024;42(2):293–304. doi:10.1038/s41587-023-01767-y

29. McKenzie AT, Katsyv I, Song WM, Wang M, Zhang B. DGCA: A comprehensive R package for Differential Gene Correlation Analysis. BMC Syst Biol. 2016;10(1):106. doi:10.1186/s12918-016-0349-1

30. Dolgalev I. MSigDB Gene Sets for Multiple Organisms in a Tidy Data Format. Published online 2026. https://igordot.github.io/msigdbr/

31. Hänzelmann S, Castelo R, Guinney J. GSVA: gene set variation analysis for microarray and RNA-Seq data. BMC Bioinformatics. 2013;14(1):7. doi:10.1186/1471-2105-14-7

32. Blighe K. EnhancedVolcano. Published online 2018. doi:10.18129/B9.BIOC.ENHANCEDVOLCANO

33. Gu Z, Gu L, Eils R, Schlesner M, Brors B. *circlize* implements and enhances circular visualization in R. Bioinformatics. 2014;30(19):2811–2812. doi:10.1093/bioinformatics/btu393

34. Jeong S, Ahn C, Kwon JS, Kim K, Jeung EB. Effects of Sodium Arsenite on the Myocardial Differentiation in Mouse Embryonic Bodies. Toxics. 2023;11(2):142. doi:10.3390/toxics11020142

35. Wu X, Chen Y, Luz A, Hu G, Tokar EJ. Cardiac Development in the Presence of Cadmium: An in Vitro Study Using Human Embryonic Stem Cells and Cardiac Organoids. Environ Health Perspect. 2022;130(11):117002. doi:10.1289/EHP11208

36. Wang Q, Ma Y, Li Y, He Z, Feng B. Lead-induced cardiomyocytes apoptosis by inhibiting gap junction intercellular communication via modulating the PKCα/Cx43 signaling pathway. Chem Biol Interact. 2023;376:110451. doi:10.1016/j.cbi.2023.110451

37. Zaffran S, Frasch M. Early Signals in Cardiac Development. Circ Res. 2002;91(6):457–469. doi:10.1161/01.RES.0000034152.74523.A8

38. de Almeida AJPO, de Oliveira JCPL, da Silva Pontes LV, et al. ROS: Basic Concepts, Sources, Cellular Signaling, and its Implications in Aging Pathways. Oxid Med Cell Longev. 2022;2022:1225578. doi:10.1155/2022/1225578

39. Zhang R, Lahens NF, Ballance HI, Hughes ME, Hogenesch JB. A circadian gene expression atlas in mammals: Implications for biology and medicine. Proc Natl Acad Sci. 2014;111(45):16219–16224. doi:10.1073/pnas.1408886111

40. Zhang YL, Li Q, Yang XM, et al. SPON2 Promotes M1-like Macrophage Recruitment and Inhibits Hepatocellular Carcinoma Metastasis by Distinct Integrin–Rho GTPase–Hippo Pathways. Cancer Res. 2018;78(9):2305–2317. doi:10.1158/0008-5472.CAN-17-2867

41. Bian ZY, Wei X, Deng S, et al. Disruption of mindin exacerbates cardiac hypertrophy and fibrosis. J Mol Med. 2012;90(8):895–910. doi:10.1007/s00109-012-0883-2

42. Giovou AE, Gladka MM, Christoffels VM. The Impact of Natriuretic Peptides on Heart Development, Homeostasis, and Disease. Cells. 2024;13(11):931. doi:10.3390/cells13110931

43. Jiménez-Ortega V, Cardinali DP, Fernández-Mateos MP, Ríos-Lugo MJ, Scacchi PA, Esquifino AI. Effect of cadmium on 24-hour pattern in expression of redox enzyme and clock genes in rat medial basal hypothalamus. BioMetals. 2010;23(2):327–337. doi:10.1007/s10534-010-9292-6

44. Hsu CY, Chuang YC, Chang FC, Chuang HY, Chiou TTY, Lee CT. Disrupted Sleep Homeostasis and Altered Expressions of Clock Genes in Rats with Chronic Lead Exposure. Toxics. 2021;9(9):217. doi:10.3390/toxics9090217

45. Sabbar M, Dkhissi-Benyahya O, Benazzouz A, Lakhdar-Ghazal N. Circadian Clock Protein Content and Daily Rhythm of Locomotor Activity Are Altered after Chronic Exposure to Lead in Rat. Front Behav Neurosci. 2017;11:178. doi:10.3389/fnbeh.2017.00178

46. Chang SJ, Chen WT, Chai CY. Arsenic-induced disruption of circadian rhythms and glutamine anaplerosis in human urothelial carcinoma. J Trace Elem Med Biol. 2024;86:127507. doi:10.1016/j.jtemb.2024.127507

47. Bourdonnay E, Morzadec C, Sparfel L, et al. Global effects of inorganic arsenic on gene expression profile in human macrophages. Mol Immunol. 2009;46(4):649–656. doi:10.1016/j.molimm.2008.08.268

48. Torres-Arellano JM, Osorio-Yáñez C, Sánchez-Peña LC, et al. Natriuretic peptides and echocardiographic parameters in Mexican children environmentally exposed to arsenic. Toxicol Appl Pharmacol. 2020;403:115164. doi:10.1016/j.taap.2020.115164

49. Wāng Y, Han Y, Xu DX. Mitochondrial metabolism reprogramming-mediated cardiomyocyte senescence involved in arsenic stress-evoked heart failure. Environ Int. 2025;202:109686. doi:10.1016/j.envint.2025.109686

50. Sundaresan S, John S, Paneerselvam G, Andiapppan R, Christopher G, Selvam GS. Gallic acid attenuates cadmium mediated cardiac hypertrophic remodelling through upregulation of Nrf2 and PECAM-1signalling in rats. Environ Toxicol Pharmacol. 2021;87:103701. doi:10.1016/j.etap.2021.103701

51. Sasikumar S, Yuvraj S, Veilumuthu P, et al. Ascorbic acid attenuates cadmium-induced myocardial hypertrophy and cardiomyocyte injury through Nrf2 signaling pathways comparable to resveratrol. 3 Biotech. 2023;13(3):108. doi:10.1007/s13205-023-03527-w

52. Liu L, Xu A, Cheung BMY. Associations Between Lead and Cadmium Exposure and Subclinical Cardiovascular Disease in U.S. Adults. Cardiovasc Toxicol. 2025;25(2):282–293. doi:10.1007/s12012-024-09955-1

53. El-Kersh K, Hopkins CD, Wu X, et al. Metallomics in pulmonary arterial hypertension patients. Pulm Circ. 2023;13(1):e12202. doi:10.1002/pul2.12202

54. Xu H, Li G, He X, et al. Low-Level Environmental Metals Exposures, Biomarkers of Myocardial Injury and Hemodynamic Stress, and Mortality Risk. JACC Adv. 2025;4(9):102064. doi:10.1016/j.jacadv.2025.102064

55. Boyett M, Li P, Xiang Y, Zhang H, Kim JK, D’Souza A. Circadian determinants of heart rhythm and arrhythmias. J Mol Cell Cardiol. 2025;208:85–101. doi:10.1016/j.yjmcc.2025.08.012

56. Shukla V, Chandrasekaran B, Tyagi A, et al. A Comprehensive Transcriptomic Analysis of Arsenic-Induced Bladder Carcinogenesis. Cells. 2022;11(15):2435. doi:10.3390/cells11152435

57. Riedmann C, Ma Y, Melikishvili M, et al. Inorganic Arsenic-induced cellular transformation is coupled with genome wide changes in chromatin structure, transcriptome and splicing patterns. BMC Genomics. 2015;16(1):212. doi:10.1186/s12864-015-1295-9

58. Ghosh P, Murumulla L, Das S, Alavala S, Challa S. Heavy metal toxicity as a driver of endoplasmic reticulum stress and dysfunction. FEBS J. Published online April 8, 2026:febs.70527. doi:10.1111/febs.70527

59. Balali-Mood M, Naseri K, Tahergorabi Z, Khazdair MR, Sadeghi M. Toxic Mechanisms of Five Heavy Metals: Mercury, Lead, Chromium, Cadmium, and Arsenic. Front Pharmacol. 2021;12:643972. doi:10.3389/fphar.2021.643972

60. Qin W, Feng J, Tian R, et al. 2-Aminoethoxydiphenyl-borate reduces arsenic-induced cardiotoxicity in rats. Acta Biochim Biophys Sin. Published online September 1, 2022. doi:10.3724/abbs.2022134

61. Shen J, Wang X, Zhou D, et al. Modelling cadmium-induced cardiotoxicity using human pluripotent stem cell-derived cardiomyocytes. J Cell Mol Med. 2018;22(9):4221–4235. doi:10.1111/jcmm.13702

62. Yang D, Yang Q, Fu N, et al. Hexavalent chromium induced heart dysfunction via Sesn2-mediated impairment of mitochondrial function and energy supply. Chemosphere. 2021;264:128547. doi:10.1016/j.chemosphere.2020.128547

63. Babiker F, Al-Kouh A, Kilarkaje N. Lead exposure induces oxidative stress, apoptosis, and attenuates protection of cardiac myocytes against ischemia–reperfusion injury. Drug Chem Toxicol. 2019;42(2):147–156. doi:10.1080/01480545.2018.1429460

64. Chou SH, Lin HC, Chen SW, et al. Cadmium exposure induces histological damage and cytotoxicity in the cardiovascular system of mice. Food Chem Toxicol. 2023;175:113740. doi:10.1016/j.fct.2023.113740

65. Souza ACF, De Paiva Coimbra JL, Ervilha LOG, et al. Arsenic induces dose-dependent structural and ultrastructural pathological remodeling in the heart of Wistar rats. Life Sci. 2020;257:118132. doi:10.1016/j.lfs.2020.118132

66. Liu Q, Xu C, Jin J, et al. Early-life exposure to lead changes cardiac development and compromises long-term cardiac function. Sci Total Environ. 2023;904:166667. doi:10.1016/j.scitotenv.2023.166667

67. Dang KD, Ho CNQ, Van HD, et al. Hexavalent Chromium Inhibited Zebrafish Embryo Development by Altering Apoptosis- and Antioxidant-Related Genes. Curr Issues Mol Biol. 2023;45(8):6916–6926. doi:10.3390/cimb45080436

68. Singh V, Singh N, Verma M, et al. Hexavalent-Chromium-Induced Oxidative Stress and the Protective Role of Antioxidants against Cellular Toxicity. Antioxidants. 2022;11(12):2375. doi:10.3390/antiox11122375

69. Wang LP, Fan SJ, Li SM, Wang XJ, Gao JL, Yang XH. Oxidative stress promotes myocardial fibrosis by upregulating KCa3.1 channel expression in AGT-REN double transgenic hypertensive mice. Pflüg Arch - Eur J Physiol. 2017;469(9):1061–1071. doi:10.1007/s00424-017-1984-0

70. Ong S, Ligons DL, Barin JG, et al. Natural Killer Cells Limit Cardiac Inflammation and Fibrosis by Halting Eosinophil Infiltration. Am J Pathol. 2015;185(3):847–861. doi:10.1016/j.ajpath.2014.11.023

71. Pan Z, Gong T, Liang P. Heavy Metal Exposure and Cardiovascular Disease. Circ Res. 2024;134(9):1160–1178. doi:10.1161/CIRCRESAHA.123.323617

72. Tohse N, Seki S, Kobayashi T, Tsutsuura M, Nagashima M, Yamada Y. Development of Excitation-Contraction Coupling in Cardiomyocytes. Jpn J Physiol. 2004;54(1):1–6. doi:10.2170/jjphysiol.54.1

73. Nakanishi T, Jarmakani JM. Developmental changes in myocardial mechanical function and subcellular organelles. Am J Physiol-Heart Circ Physiol. 1984;246(4):H615–H625. doi:10.1152/ajpheart.1984.246.4.H615

74. Aballo TJ, Bae J, Paltzer WG, et al. Integrated proteomics identifies troponin I isoform switch as a regulator of a sarcomere-metabolism axis during cardiac regeneration. Cardiovasc Res. 2025;121(8):1240–1253. doi:10.1093/cvr/cvaf069

75. Cao T, Liccardo D, LaCanna R, et al. Fatty Acid Oxidation Promotes Cardiomyocyte Proliferation Rate but Does Not Change Cardiomyocyte Number in Infant Mice. Front Cell Dev Biol. 2019;7:42. doi:10.3389/fcell.2019.00042

76. Gogiraju R, Bochenek ML, Schäfer K. Angiogenic Endothelial Cell Signaling in Cardiac Hypertrophy and Heart Failure. Front Cardiovasc Med. 2019;6:20. doi:10.3389/fcvm.2019.00020

77. Giudice J, Xia Z, Wang ET, et al. Alternative splicing regulates vesicular trafficking genes in cardiomyocytes during postnatal heart development. Nat Commun. 2014;5(1):3603. doi:10.1038/ncomms4603

78. Rein KA, Borrebaek B, Bremer J. Arsenite inhibits β-oxidation in isolated rat liver mitochondria. Biochim Biophys Acta BBA - Lipids Lipid Metab. 1979;574(3):487–494. doi:10.1016/0005-2760(79)90245-5

79. Vaziri ND. Mechanisms of lead-induced hypertension and cardiovascular disease. Am J Physiol-Heart Circ Physiol. 2008;295(2):H454–H465. doi:10.1152/ajpheart.00158.2008

80. Zhou X, Hao W, Shi H, Hou Y, Xu Q. Calcium Homeostasis Disruption - a Bridge Connecting Cadmium-Induced Apoptosis, Autophagy and Tumorigenesis. Oncol Res Treat. 2015;38(6):311–315. doi:10.1159/000431032

81. Thévenod F, Lee WK. Cadmium and cellular signaling cascades: interactions between cell death and survival pathways. Arch Toxicol. 2013;87(10):1743–1786. doi:10.1007/s00204-013-1110-9

82. Bianchetti L, Barczyk M, Cardoso J, Schmidt M, Bellini A, Mattoli S. Extracellular matrix remodelling properties of human fibrocytes. J Cell Mol Med. 2012;16(3):483–495. doi:10.1111/j.1582-4934.2011.01344.x

83. Bunger MK, Wilsbacher LD, Moran SM, et al. Mop3 Is an Essential Component of the Master Circadian Pacemaker in Mammals. Cell. 2000;103(7):1009–1017. doi:10.1016/S0092-8674(00)00205-1

84. Lopez-Molina L, Conquet F, Dubois-Dauphin M, Schibler U. The DBP gene is expressed according to a circadian rhythm in the suprachiasmatic nucleus and influences circadian behavior. EMBO J. 1997;16(22):6762–6771. doi:10.1093/emboj/16.22.6762

85. Martínez-González J, Cañes L, Alonso J, et al. NR4A3: A Key Nuclear Receptor in Vascular Biology, Cardiovascular Remodeling, and Beyond. Int J Mol Sci. 2021;22(21):11371. doi:10.3390/ijms222111371

86. Lu H, Delnicki M, Griffin G, Wise JL. Current Understanding of Sex Differences in Metal-Induced Diseases. Curr Environ Health Rep. 2025;12(1):18. doi:10.1007/s40572-025-00482-x

87. Wang Z, Sun Y, Yao W, Ba Q, Wang H. Effects of Cadmium Exposure on the Immune System and Immunoregulation. Front Immunol. 2021;12:695484. doi:10.3389/fimmu.2021.695484

88. Zhang H, Wang J, Zhang K, et al. Association between heavy metals exposure and persistent infections: the mediating role of immune function. Front Public Health. 2024;12:1367644. doi:10.3389/fpubh.2024.1367644

89. Balabekova MK, Kairanbayeva GK, Yu VK, et al. Immunomodulatory Effects of a New Ethynylpiperidine Derivative: Enhancement of CD4+FoxP3+ Regulatory T Cells in Experimental Acute Lung Injury. Biomedicines. 2025;13(12):3017. doi:10.3390/biomedicines13123017

